# Identifying the function of the NMDA NR1^C2^ subunit through its interaction with Magi-2 during inflammatory pain

**DOI:** 10.1101/2024.01.30.578033

**Authors:** Garrett D. Sheehan, Molly K. Martin, Adam Roszczyk, Katherient Hao, Arin Bhattacharjee

**Affiliations:** Program for Neuroscience, University at Buffalo - The State University of New York, Buffalo, New York 14203, USA; Pharmacology and Toxicology, University at Buffalo - The State University of New York, Buffalo, New York 14203, USA

## Abstract

Much is understood about the structure and gating properties of NMDA receptors (NMDAR), but the function of the carboxy-terminal splice variant of the NR1 subunit, NR1^C2^ has never been identified. By studying the scaffolding protein Magi-2 in animal models of inflammatory pain, we discovered how NR1^C2^ protein is specifically regulated. We found that Magi-2 deficiency resulted in decreased pain behavior and a concomitant reduction in NR1^C2^ protein. Magi-2 contains WW domains, domains typically found in ubiquitin ligases. We identified an atypical WW-binding domain within NR1^C2^ which conferred susceptibility to Nedd4-1 ubiquitin-ligase dependent degradation. We used lipidated peptidomimetics derived from the NR1^C2^ sequence and found that NR1^C2^ protein levels and pain behavior can be pharmacologically targeted. The function of NR1^C2^ is to give lability to a pool of NMDAR, important for pain signaling.

## Introduction

NMDA receptors (NMDAR) are essential for pain transmission ^1^. In 1992, Coderre and Melzack made a seminal observation: intrathecal administration of selective NMDA receptor antagonists, but not AMPA receptor antagonists, blocked inflammatory pain behavior in mice ^2^. NMDA receptor antagonists have been used clinically to treat various pain conditions with mixed results. For example, ketamine provides analgesic and antihyperalgesic effects, however, pain relief is modest, suggesting that ketamine is only useful when opioids are contraindicated ^3^. Ketamine is not selective for specific NMDAR subtypes ^4^ and therefore at increasing doses, there will be many adverse off-target effects. This limits the use of current NMDAR antagonists at higher concentrations, which might be required for effective pain treatment. A better understanding of NMDAR activity, and importantly a more specific NMDAR antagonism approach may improve clinical success.

A functional NMDAR is a heterotetramer composed of two obligate NR1 subunits and two NR2 (and in some cases NR3 subunits) ^5^. There are 8 splice isoforms of NR1, each of which are expressed in the spinal cord ^6,7^. The last stretch of the NR1 C-terminal domain consists of the C2 cassette. Alternative splicing of the C2 cassette produces two distinct C-termini called C2 and C2’. The C2’ isoform contains a PDZ binding motif shown to bind to PSD-95 and associated postsynaptic density proteins ^8^. The C2 isoform does not contain a PDZ binding motif, and the function has yet to be resolved. Functional NMDA receptors containing only the NR1^C2’^ subunit will have a total of 4 PDZ binding motifs, as each of the two NR2 subunits also possess a PDZ binding motif. By extension, NMDAR containing only the NR1^C2^ subunit will have 2 PDZ binding motifs, half the number found in NMDAR with NR1^C2’^, presumably reducing their affinity for the postsynaptic density. Magi-2 is a scaffold protein which was first discovered by its ability to bind to NMDAR ^9^. Like PSD-95, Magi-2 contains multiple PDZ binding domains and a catalytically inactive guanylate kinase-like (GUK) domain. Unlike PSD-95, Magi proteins additionally contain two WW domains ^10^. WW-binding domains are typically found in ubiquitin ligases and are used to recognize substrates for ubiquitination and subsequent degradation ^11^. The sequence of the first WW domain of Magi-2 is notably like the WW domains of the ubiquitin ligase Nedd4-1 ^9^. By competing with ubiquitin ligases for substrate recognition, Magi protein-WW domains confer substrate protection from ubiquitin ligase-dependent degradation ^12–14^.

Protein ubiquitination acts as a signal for the sorting, trafficking, and endocytosis of membrane proteins. Multiple ubiquitin ligases are known to specifically regulate the functions of ion channels, transporters, and signaling receptors ^15^. The Nedd4 E3 ubiquitin ligases specifically translocate to the membrane in response to calcium ^16^. Furthermore, dendritic proteasomal function is essential for synaptic homeostasis and plasticity ^17^. Here we identify a Group III WW-binding domain comprising a proline-arginine motif (-L/PPR-) ^18–20^ in NR1^C2^. We show how this site is important for Nedd4-1-dependent degradation of NR1^C2^. We demonstrate that Magi-2 deficiency results in reduced NR1^C2^ levels which attenuates inflammatory pain behaviors. Using lipidated peptidomimetics derived from the NR1^C2^ sequence we demonstrate that NMDAR trafficking can be pharmacologically modulated.

## Results

### Magi-2 deficiency is associated with NMDAR hypofunctioning

We were initially interested in the putative role of Magi-2 in inflammatory pain signaling (Fig. 1A). The spinal cord dorsal horn (SCDH) is canonically subdivided into anatomically defined laminae, delineated by the class of sensory neurons that terminate within them ^21^. We first performed Magi-2 immunohistochemistry using a monoclonal antibody against Magi-2 comparing the expression pattern to the molecular marker PKC*γ*, which is found exclusively at the border of lamina 2i and 3o (Fig. 1B). This antibody was validated by immunocytochemistry during heterologous expression of Magi-2 (Fig. S2). Magi-2 expression was observed superficial to the band formed by PKC*γ* immunoreactivity (Fig. 1B) suggesting Magi-2 is present in second order neurons receiving input from high threshold nociceptors including C-fibers ^22^. This finding was recapitulated using a previously validated polyclonal antibody against Magi-2, where both immunohistochemistry and Western analysis confirmed Magi-2 expression within the SCDH (Figs S1&S3). We subsequently acquired a Magi-2 floxed mouse line (Fig S4A, B) which had previously been used to study Magi-2 in kidney podocyte formation ^13^. Magi-2 protein analysis however, revealed that the homozygous floxed mice (hereafter referred to as ΔMagi-2) had endogenously reduced levels of Magi-2 alpha and beta isoforms when compared to wildtype (WT) and heterozygotic littermates and were devoid of the gamma isoform (Fig 1C & D, Fig S4C, D). We understood that performing Cre-mediated, targeted deletion of Magi-2 using this mouse line would be confounding. Nonetheless, we used this endogenous global Magi-2 deficiency to test for potential pain behavior phenotypes. Baseline evoked thermal (Hargreaves) and light touch (von Frey) responses were normal across genotypes (Fig S4E, F) however, ΔMagi2 mice lacked formalin-induced nocifensive behaviors (licks and lifts directed at the injected paw) associated with the second, inflammatory phase of this assay (Fig 1E&F).

**Figure 1.**
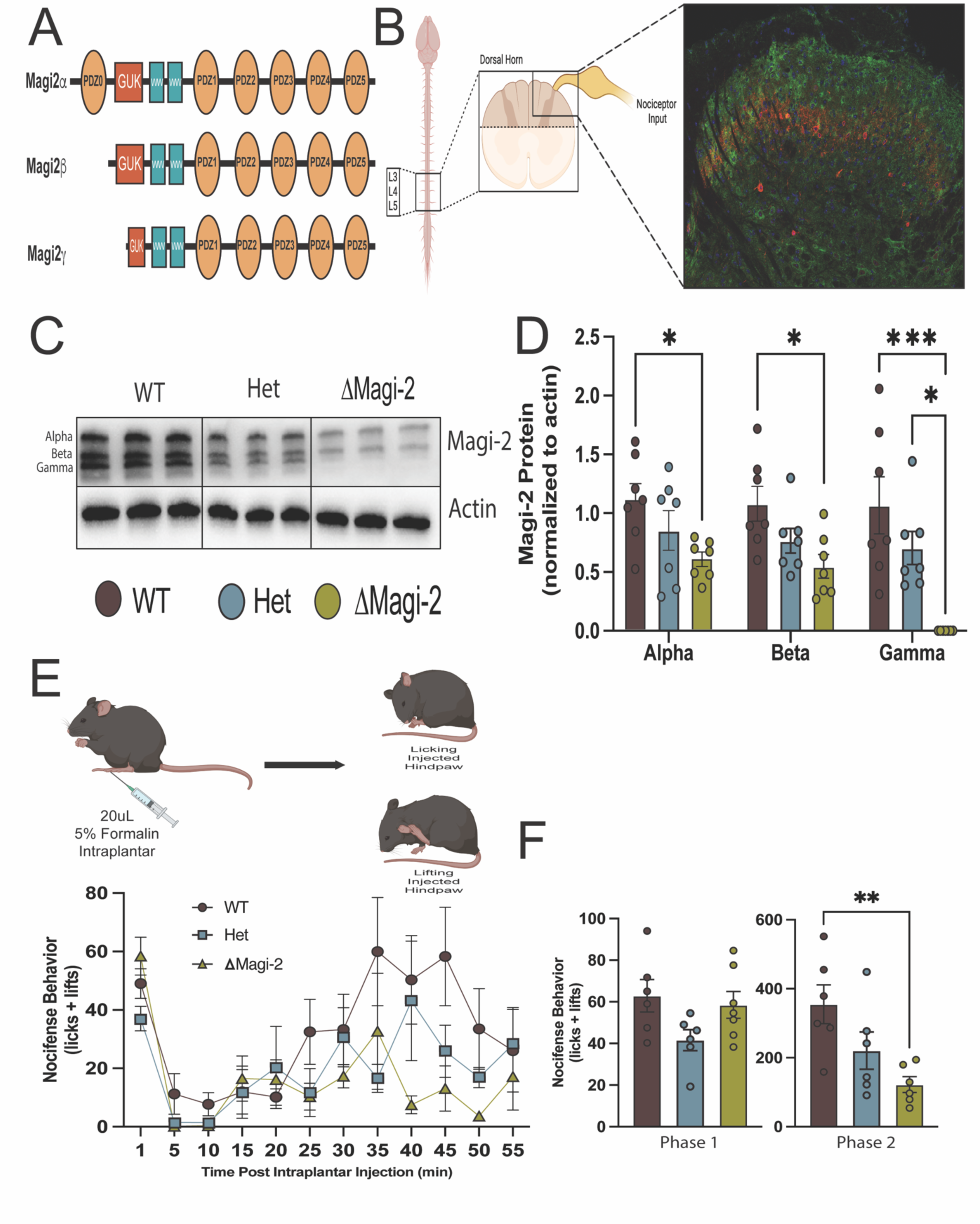
A Magi-2 floxed line has endogenous Magi-2 deficiency and exhibits diminished inflammatory pain behavior. **A)** Schematic of the three isoforms of Magi-2. The alpha, beta, and gamma isoforms differ with respect to whether they contain the PDZ-0 and GK domains. **B)** Immunolabeling of Magi-2 (green) in the lumbar spinal cord dorsal horn revealed superficial expression relative to the lamina 2/3 border as defined by PKC*γ* (red) expression. **C**) Western analysis of Magi-2 protein levels from the synaptic fraction of dorsal horn lysates of WT, heterozygous and homozygous (floxed) ΔMagi-2 mice. ΔMagi-2 mice exhibit reduced Magi-2 alpha and beta isoforms expression and no Magi-2 gamma expression. Actin was used as a loading control. Individual lanes represent one replicate from a single lysate which run in triplicate. **D**) Densitometric quantification of relative Magi-2 protein levels normalized to actin from 1C. Each set of three bands from a single lysate is averaged to create an n of 1, for a total of n = 7 different mice for each genotype. Relative protein levels of each isoform were compared by One-way ANOVA with Tukey post hoc test. (Magi-2 Alpha: WT vs. ΔMagi-2, p= 0.0369*; Magi-2 Beta: WT vs ΔMagi-2, p=0.0146*; Magi-2 Gamma: WT vs ΔMagi-2, p=0.0006***, Het vs ΔMagi-2 p=0.0173*). **E**) ΔMagi-2 mice exhibit deficits in the formalin assay of inflammatory pain. Mice received a unilateral 20*μ*L intraplantar injection of 5% formalin and subsequent licking and lifting behaviors were scored by a blinded observer. Nocifensive behaviors (defined as the sum of lifts and licks directed at the injected hindpaw of mice injected with formalin and video recorded for one hour. **F)** Cumulative nocifensive behaviors for Phase 1 (0-5min) and Phase 2 (10-55min) following intraplantar injection are shown in the bar graphs. (WT vs ΔMagi-2, p=0.0085** One-Way ANOVA with Tukey post hoc test).

Multiple studies have shown the involvement of SCDH NMDAR as necessary for formalin-induced inflammatory pain behaviors ^23–25^. Consequently, we probed for any alterations in NMDAR levels in the crude synaptosomal fraction of SCDH lysates collected from WT, heterozygous, and ΔMagi-2 mice (Fig 2A, B). GluN2A has been shown to be the dominant isoform in the adult central nervous system ^26^ and GluN2B is the isoform primarily responsible for synaptic transmission in lamina 1 ^27^. However, we found no changes in either NR2A or NR2B when comparing lysates from WT, heterozygous or ΔMagi-2 mice (Fig 2A&B). Conditional deletion of the NR1 subunit in the SCDH, while not altering basal pain behavior, reduced formalin-induced pain^28^. Other studies have also shown a reduction in nocifensive behaviors in the second phase of the formalin assay during targeted deletion of NR1 in the dorsal horn ^29–31^. In WT mice, we co-immunoprecipitated NR1 with Magi-2 from dorsal horn lysates (Fig. 2C), replicating prior findings from the brain ^9^. In ΔMagi-2 mice, we also found a gene-copy number-dependent reduction in NR1 protein levels (Fig 2B). With this approach, we were unable to distinguish between NMDAR located postsynaptically in dorsal horn neurons from those located on the presynaptic terminals of primary afferents ^32^. We therefore performed whole-cell voltage-clamp recordings on second order neurons in lamina 1of the SCDH to determine whether the reduction in NR1 protein levels affected postsynaptic NMDA-evoked current densities. A recent study showed that the synaptic pool of NMDAR was unaffected by a genetic loss of NR1 due to a compensatory shift in NMDAR from extrasynaptic to synaptic sites ^33^. Consequently, we chose to bath apply N-Methyl-D-Aspartate (NMDA) followed by washout and found an overall reduction in NMDA-induced currents in dorsal horn neurons (Fig 2D-G). In contrast, we observed no differences in spontaneous excitatory postsynaptic current amplitude, frequency, nor Tau of decay of averaged events and saw no protein level changes in either of the AMPA receptor (AMPAR) subunits GluA1 or GluA2 (FigS5). Results from the ΔMagi-2 mouse strain suggested that Magi-2 deficiency affected inflammatory pain through its regulation of NR1 subunit protein levels.

**Figure 2.**
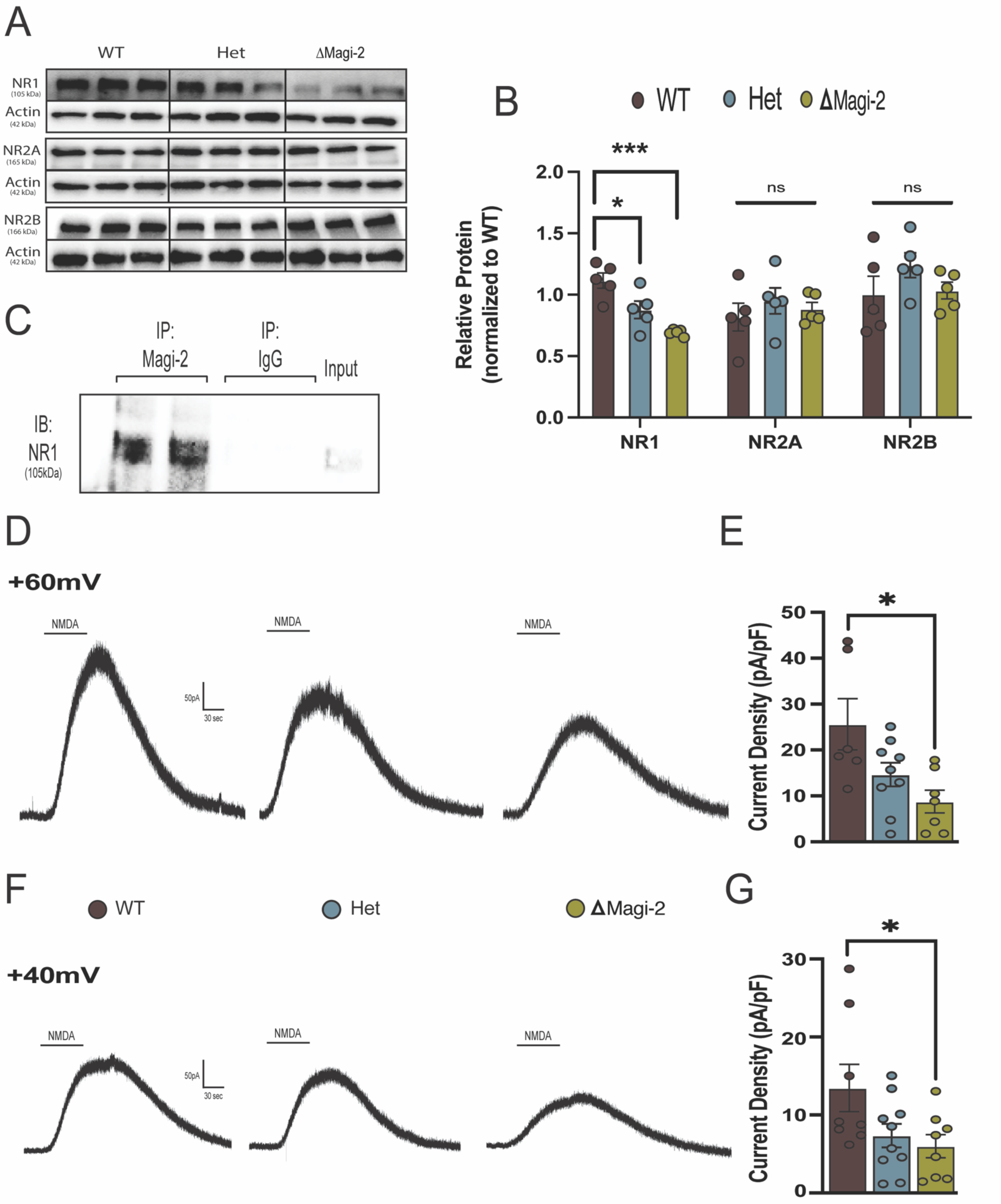
Magi-2 deficiency is associated with a decrease in NR1 protein and decreased NMDA currents in the SCDH. **A**) Western analysis of NR1, NR2A, and NR2B NMDA receptor levels from spinal cord dorsal horn lysates of WT, heterozygous and homozygous ΔMagi-2 mice with actin as a loading control. **B**) Densitometric analysis of 2A. Relative protein levels of each NMDA receptor subunit were compared using one-way ANOVA with Tukey post hoc analysis (**NR1**: WT vs. Het p=0.0249*; WT vs ΔMagi-2 p = 0.0004***) **C)** Immunoblot of NR1 following Magi-2 co-immunoprecipitation from dorsal horn lysates. **D & F)** Representative NMDA-evoked current traces recorded from lamina 1 neurons of WT, heterozygous, and ΔMagi-2 mice. Neurons were held at **D)** +60mV and **F**) +40mV and then 50uM NMDA was bath applied for 60 seconds followed by a washout for 3 minutes. **E & G)** Current densities (peak NMDA-evoked current amplitude divided by cell capacitance compared by One-way ANOVA with Tukey post-hoc test. **E**) +60mV: WT vs ΔMagi-2 p= 0.0392* **G**) +40mV: WT vs. ΔMagi-2 p=0.0436*: one-way ANOVA with Tukey post hoc test.

### Magi-2 knockdown in wildtype mice regulates NR1^C2^ protein expression and decreases pain behavior

To avoid any confounding developmental effects present in the ΔMagi-2 mouse line we decided to use a viral Magi-2-shRNA knockdown approach in wildtype mice. AAV2/9-GFP-U6-m-Magi-2-shRNA was administered intrathecally into the lumbar dorsal horn of male and female WT C57Bl/6 mice (Fig 3A) to assess the effect of Magi-2 knockdown in the Complete Freunds’ Adjuvant (CFA) model of chronic inflammatory pain. The Magi-2 shRNA sequence was previously validated ^34^. We first confirmed viral transduction of the dorsal horn via immunohistochemical staining for GFP (Fig S6). Intrathecal injection of Magi-2-shRNA led to a reduction in all three isoforms of Magi-2 as measured by Western analysis from lumbar SCDH lysates collected 2 weeks following injection (Fig 3B°C; Fig S6). In the CFA inflammatory pain assay, Magi-2 knockdown mice exhibited increased thermal paw withdrawal latencies (PWL) compared to scrambled control (AAV2/9-GFP-U6-m-scram) treated mice (Fig. 3D). Baseline PWL prior to CFA injection was unaffected by Magi-2 knockdown (Fig. S7). Both male and female mice were tested in this experiment and while slight differences in the dynamics of CFA-induced thermal hyperalgesia were observed, there were no direct sex-dependent differences in PWL (Fig S7). Using the Dynamic Weight Bearing assessment, we observed that mice injected with the scrambled shRNA exhibited a significant reduction in the percentage of time spent placing weight on the CFA injected hindpaw, but this decrease was absent in Magi-2 knockdown mice(Fig 3E&F).

**Figure 3.**
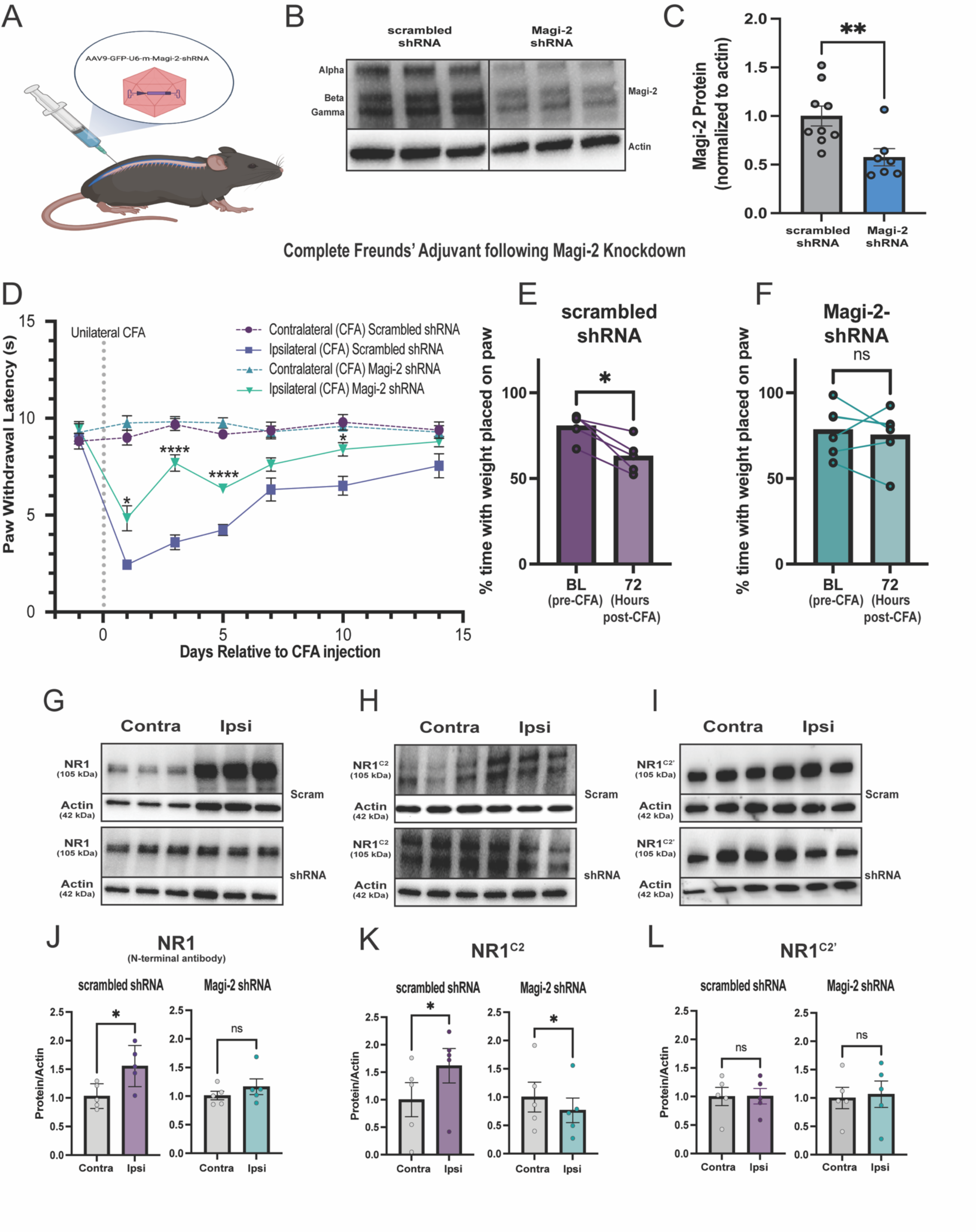
Viral knockdown of Magi-2 in wildtype mice results in reduced chronic inflammatory pain behavior and a specific decrease in NR1^C2^ protein. **A**) Mice were injected intrathecally with either AAV2/9-GFP-U6-m-Magi-2-scrambled or AAV2/9-GFP-U6-m-Magi-2-shRNA. **B**) Western analysis of total Magi-2 protein levels from spinal cord dorsal horn of mice two weeks after receiving either AAV2/9-GFP-U6-m-Magi-2-scrambled or AAV2/9-GFP-U6-m-Magi-2-shRNA relative to the actin loading control. **C**) Densitometric analysis of 3B compared using unpaired student’s t-test: scrambled shRNA vs Magi-2 shRNA p=0.0090**. Sum of all isoforms were included in the analysis. **D**) Thermal withdrawal latencies following unilateral intraplantar injection of CFA 2 weeks after intrathecal injection of either of AAV2/9-GFP-U6-m-Magi-2-shRNA or scrambled shRNA control. Baseline responses were recorded two days prior to AAV administration and for two weeks until CFA injection. Two-way ANOVA with Bonferroni correction for paw withdrawal latencies following CFA injection (ipsilateral scrambled shRNA vs ipsilateral Magi-2 shRNA: 24 hours: p=0.0233*; 3 days: p<0.0001****; 5 days: p<0.0001****; 10 days: p=0.0308* **E**) Scrambled shRNA injected mice spend less time on the hind paw after injection with CFA as measured via Dynamic Weight Bearing. Student’s paired t-test (scrambled shRNA: Baseline vs. 72 hours post CFA p = 0.0104*). **F**) Magi-2 shRNA injected mice spend an equivalent amount time spent on the hind paw after injection of CFA compared to baseline as measured via Dynamic Weight Bearing. No significant change from baseline was observed **G**) Western analysis of total NR1 protein levels in the ipsilateral vs contralateral dorsal horn following unilateral intraplantar injection of CFA (72 hours) two weeks following intrathecal Magi-2 knockdown using an N-terminal targeting antibody which recognizes both C-terminal splice isoforms. In the scrambled group, there was an increase in NR1 expression. **H**) Western analysis of NR1^C2^ levels from the same lysates as G/H. **I)** Western analysis of NR1^C2’^ levels from the same lysates as G/H. **J)** Densitometric quantification of **G**, paired students t-test (scrambled: Ipsilateral vs. contralateral p=0.0231*) **K)** Densitometric quantification of **I**: paired t-test: scrambled: ipsilateral vs. contralateral p=0.0216*; Magi-2 shRNA: ipsilateral vs. contralateral p=0.0178*. **L**) Densitometric quantification of **I**

We collected dorsal horn lysates 48 hours after unilateral intraplantar CFA injection from mice receiving either scrambled control or Magi-2-shRNA intrathecally two weeks prior. Control mice showed an increase in NR1 levels in the dorsal horn ipsilateral to the injected paw, like in prior studies^35^. In contrast, mice that had received Magi-2 shRNA showed no difference in dorsal horn NR1 levels when comparing ipsilateral and contralateral sides. (Fig 3 G& J). We next sought to determine if this increase in NR1 was specific to either of the C-terminal splice isoforms of NR1. Using the same lysates, we probed for NR1^C2^ and NR1^C2’^ levels using isoform specific antibodies ^36–38^. NR1^C2’^ levels were unchanged in either the scrambled shRNA condition or the Magi-2 shRNA condition. On the other hand, NR1^C2^ was specifically increased in the ipsilateral dorsal horn of mice that received scrambled shRNA and there was a significant decrease in the mice that had received Magi-2 shRNA (Fig 3H,K & I,L). These results suggests that the increase in NR1 seen in CFA is specific to the NR1^C2^ splice isoform and this increase is dependent on the presence of Magi-2. To further address the specific effect of Magi-2 on NR1^C2^ levels, we re-examined the lysates from the ΔMagi-2 SCDH lysates and found that the loss of NR1 was specific to the NR1^C2^ isoform, with no differences in NR1^C2’^ (Fig S8) further supporting a role for Magi-2 in specifically regulating NR1^C2^ protein levels.

### A WW binding domain within NR1^C2^ confers sensitivity to Nedd4-1-dependent degradation

Our results showed that overall NR1^C2^ abundance is dependent upon Magi-2 expression. We therefore hypothesized that NR1^C2^ is likely regulated by ubiquitin ligase activity. The WW domain-containing E3 ubiquitin ligase Nedd4-1 canonically recognizes substrates containing Group I, (-L/PPXY-) WW binding domains, with X referring to any amino acid ^39^. Group III WW binding domains containing a proline-arginine motif (-L/PPR-) have also been characterized ^18–20^. Within the C-terminal sequence of the NR1^C2^ variant, we identified an -LPR-motif (Fig 4A). Interestingly, NR1^C2^ is only found in mammals, and the -LPR-motif is highly conserved within mammalian species (Fig. S9). We hypothesized that the -LPR-motif allows regulation by Nedd4-1 (Fig 4A). To test this, we co-transfected CHO cells with NR1^C2^ and Nedd4-1 and observed a significant reduction in NR1^C2^ protein levels (Fig 4B, C). This loss of protein was rescued by treatment of transfected cells with the proteasome inhibitor MG-132 (Fig S10), suggesting that ubiquitin-dependent proteasomal degradation was responsible for the loss of NR1^C2^ protein in this assay. To demonstrate that this specific sequence motif bestows susceptibility to ubiquitin-dependent degradation, we mutated the proline within this motif to an alanine (Fig 4A). The NR1^C2^ – P918A mutation prevented Nedd4-1 dependent degradation of NR1^C2^ (Fig 4B, D). Moreover, degradation was specific to the C2 variant as Nedd4-1 was unable to degrade NR1^C2’^ protein (Fig 4B, E).

**Figure 4.**
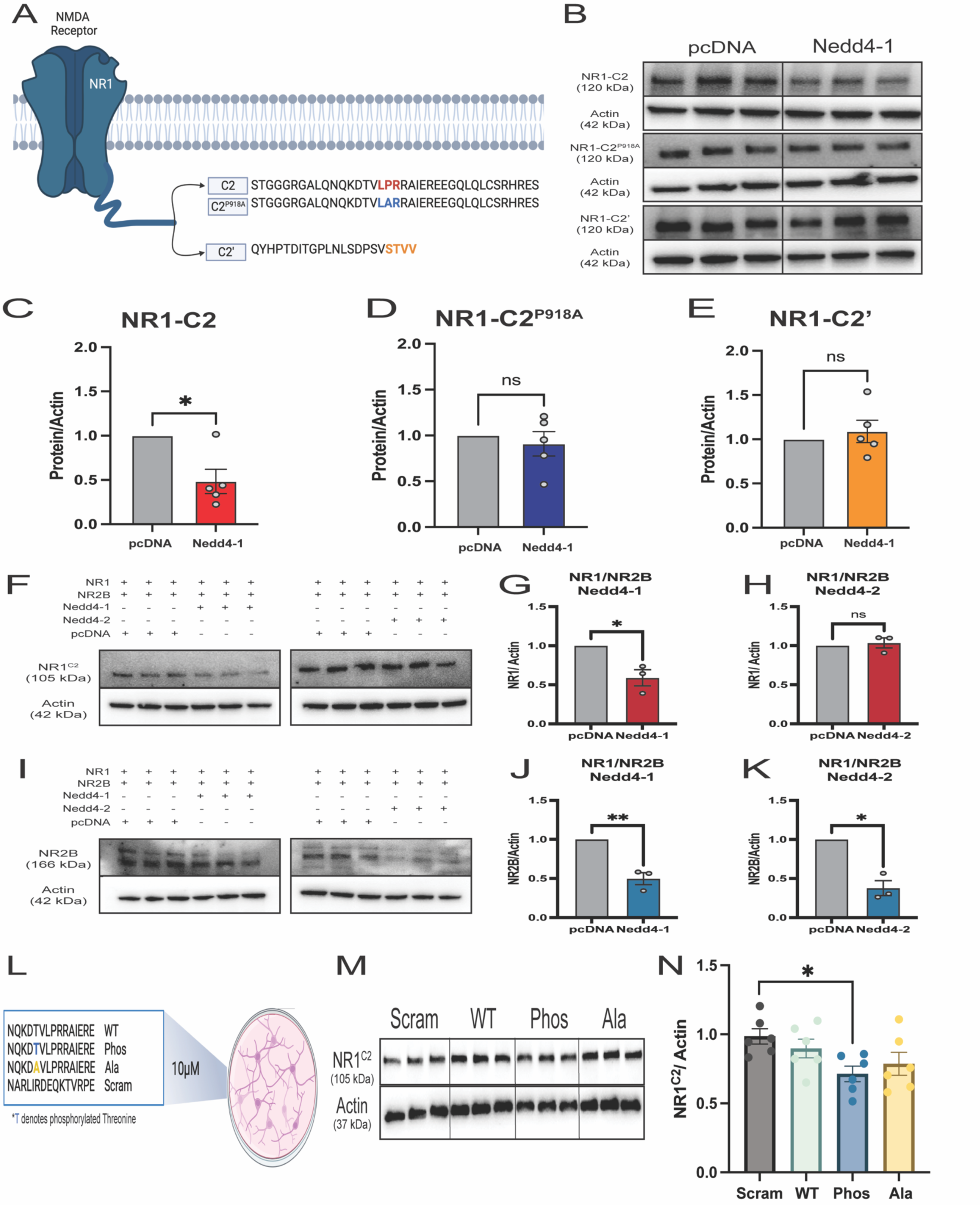
NR1^C2^ is a substrate of the Nedd4-1 ubiquitin ligase. **A)** Schematic of the NR1^C2^ and NR1^C2’^ carboxy-terminal splice isoforms of NR1 as well as the engineered mutation of the NR1^C2^ Group 3 WW-binding domain construct. **B**) Western analysis depicting the effect of co-transfection of NR1^C2^, NR1^C2’^ or NR1^C2^ ^P918A^ with the E3 ubiquitin ligase Nedd4-1 compared to pcDNA control. **C-E**) Densitometric Analysis for **B**: **(C)** NR1^C2^: pcDNA vs Nedd4-1 p = 0.0201* n =5 per group, paired students t-test **F)** Western analysis of NR1^C2^ coexpressed with NR2B to demonstrate the effect of Nedd4-1(left) or Nedd4-2 (right) on presumably fully formed channels. **G)** Densitometric analysis shows a reduction in NR1 when coexpressed with NR2B and Nedd4-1 (pcDNA vs. Nedd4-1 p = 0.0417*: paired students t-test) H) No difference was observed in NR1 levels when co-transfected with NR2B and Nedd4-2. **I)** Western analysis of NR2B coexpressed with NR1^C2^ to demonstrate the effect of Nedd4-1(left) of Nedd4-2 (right) presumably fully formed channels. **J)** Densitometric analysis shows a reduction in NR2B when coexpressed with NR1^C2^ and Nedd4-1 (pcDNA vs. Nedd4-1 p = 0.0048**: paired students t-test) **K)** Densitometric analysis shows a reduction in NR2B when coexpressed with NR1^C2^ and Nedd4-2 (pcDNA vs. Nedd4-2 p = 0.0160*: paired students t-test) **L)** NR1^C2^-based mimetic peptides corresponding to the Group 3 WW-binding domain. Unstimulated cultured hippocampal neurons were treated with 10uM peptide overnight after 7 DIV. **M)** Western analysis of NR1^C2^ levels following overnight treatment of 10uM peptide. **N)** Densitometric Analysis of **M** shows a reduction in NR1^C2^ protein levels upon treatment with the phosphorylated peptide compared to scrambled control peptide (scrambled vs. Phosphorylated peptide p=0.0480*: one-way ANOVA with Tukey post hoc test, n=6/peptide)

We further examined if Nedd4-1 acts on the NR2 subunits, NR2A and NR2B. Neither NR2A nor NR2B protein levels were reduced when co-expressed in CHO cells with Nedd4-1. NR2B, however, was reduced in the presence of Nedd4-2, an E3-ubiquitin ligase of the same family (Fig. S11). Interestingly, in fully formed NMDAR, when NR1^C2^ and NR2B are co-expressed, both NR1^C2^ and NR2B displayed sensitivity to Nedd4-1. Yet, in the presence of Nedd4-2, NR2B was still degraded but NR1^C2^ remained insensitive to Nedd4-2. These results suggest that NR1^C2^ specifically confers Nedd4-1 susceptibility on fully formed NMDAR *in vitro* (Fig 4F-L).

### NR1^C2^-based peptidomimetics bidirectionally regulate pain behavior

Our results suggest that a competition for the NR1^C2^ WW-binding domain between the WW domains of Nedd4-1 and Magi-2 exists to regulate NR1^C2^ levels and membrane expression. Consequently, we developed several peptides derived from the C-terminal amino acid sequence of NR1^C2^, derived specifically from the region immediately surrounding, and including the WW-binding domain (Fig 4F). Peptides were myristoylated at the N-termini to make them cell-penetrable and membrane delimited. After confirming the presence of Magi-2 in cultured hippocampal neurons (Fig S12), we treated them with 10uM peptide. 24 hours after treatment, we assessed the total NR1^C2^ levels by Western analysis in unstimulated neurons. The sequence based on the wildtype NR1^C2^ did not affect NR1^C2^ levels, but our prior work has shown that the phosphorylation state of mimetic peptides targeting the ubiquitination of membrane proteins can have opposing effects on protein levels^40^. Threonine 915 is the only phosphorylatable residue within the C2 sequence near the WW binding domain (Fig 4F). We reasoned that the threonine at position 915 in the NR1^C2^ carboxy terminal is phosphorylated to either promote or prevent ubiquitin ligase interaction ^41^. We therefore used two other mimetic peptides: one with the threonine phosphorylated and the other with threonine substituted with alanine to prevent potential endogenous phosphorylation of the peptide (Fig 4F). Only the phosphorylated threonine peptide caused a significant reduction in NR1^C2^ protein levels (Fig 4H). This suggests that the phosphorylation status of NR1^C2^ determines its interaction with either Magi-2 or Nedd4-1. We further explored the potential of these peptides to produce analgesia during inflammatory pain.

We intrathecally administered NR1^C2^ WW-binding domain mimetic peptides after established CFA inflammatory pain in mice (Fig 5A). Thermal PWL of male and female mice were first measured for two days prior to CFA injection into the right paw. At 24 hours after CFA injection, PWL was measured to validate sensitization. Immediately following PWL assessments, mimetic peptides and the scrambled control peptide were intrathecally injected and PWL was measured at various timepoints after injection. The wildtype NR1^C2^ sequence peptide did not affect pain behavior, however, intrathecal injection of the phosphorylated peptide rapidly reversed CFA-induced inflammatory pain in both males and females (Fig. 5B-C). This was consistent with the effect seen on NR1^C2^ levels in hippocampal cultures (Fig 4F-H). The phosphorylated peptide reduced hyperalgesia to baseline for the duration of the thermal sensitivity assay and attenuated dynamic weight bearing deficits (Fig 5G). Surprisingly, the alanine-substituted peptide prolonged thermal hyperalgesia, preventing the natural recovery observed in scrambled controls (Fig5B-D). Contralateral measurements revealed no differences amongst the peptides demonstrating that peptides exert their effect are specifically during increased SCDH activity due to pain (Fig. S13). These results suggest that these peptides possess the ability to bidirectionally control pain behavior by targeting the interaction of NR1^C2^ with either Magi-2 or Nedd4-ubiquitin ligases.

**Figure 5.**
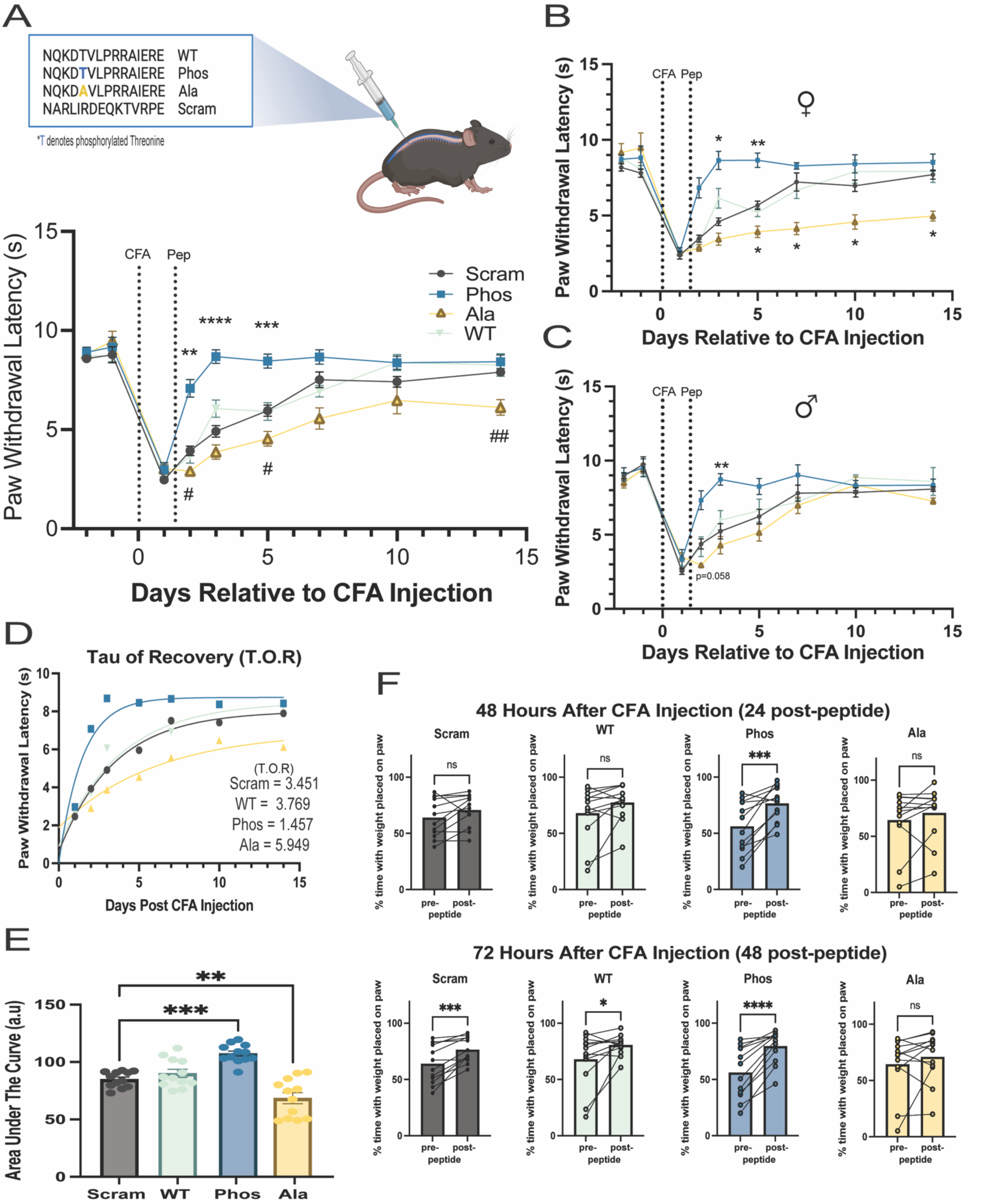
Bidirectional control of pain behavior using NR1^C2^-derived peptidomimetics. **A)** Upper: Schematic depicting intrathecal injection of NR1 lipidated peptidomimetics into WT C57Bl/6 mice 24 hours following unilateral intraplantar injection of CFA. Lower: Following two days of baseline measurements, thermal withdrawal latencies were measured 24 hours after unilateral intraplantar injection of CFA. Mice were then immediately injected intrathecally with the NR1-targeted peptidomimetics, and thermal withdrawal latency was measured 48 hours, 72 hours, and 5, 7, 10, and 14 days after CFA. Paw withdrawal latencies for Hargreaves Thermal assay for mice receiving NR1-targeted peptides. n=12 mice per peptide (6 males, 6 females). Statistically significant differences in mean paw withdrawal latency (sec) +/- SEM are given for scrambled vs Phosphorylated peptide (48 hours post CFA p=0.0012**; 72 hours p<0.0001****; 5 days p=0.0009***) and Scrambled vs Alanine peptide (48 hours p=0.0201^#^; 5 days p=0.0313^#^; 14 Days p=0.0045^##^) two-way ANOVA with Bonferroni Correction **B**) Females from **A.** Scrambled vs Phosphorylated Peptide (72 hours p=0.0128*; 5 days p=0.0044**) and Scrambled vs Alanine (5 days p=0.019^#^; 7 days p=0.0122^#^; 10 days p=0.0357^#^; 14 days p=0.0134^#^). **C**) Males from **A**: Scrambled vs Phosphorylated Peptide (72 hours p=0.0084**) two-way ANOVA with Bonferroni correction **D**) Tau analysis for recovery from CFA with lines of best fit demonstrating the rates of recovery from CFA-induced hyperalgesia. A one-phase decay line was fit to the data and the tau of recovery was calculated. Tau values are as follows: scrambled t=3.451; phosphorylated t=1.457; alanine t=5.949; WT t=3.769 **E**) Area under the curve for Thermal withdrawal latencies shows a significant difference between scrambled vs phosphorylated peptide (p=0.0003***) and scrambled vs alanine (p=0.0050**) one way ANOVA with Tukey post hoc test **F**) Time spent bearing weight on the CFA-injected hindpaw as measured via Dynamic Weight Bearing. **(Top)** At 48 hours following CFA injection (24 hours post-peptide) only the group receiving the phosphorylated peptide showed a significant increase in the amount of time spent placing weight on the injected hindpaw indicative of recovery (Phosphorylated: pre-peptide injection vs post-peptide injection p=0.001***). **(Bottom)** At 72 hours, only the mice receiving alanine peptide did not display any increase in time spent placing weight on the CFA-injected paw, while all other groups trended toward recovery (scrambled: pre- vs post-peptide injection p=0.0009***; phosphorylated: pre- vs post-peptide injection (24 vs. 72) p<0.0001****; WT: pre- vs post- peptide injection (24 vs. 72) p=0.0470*) paired students t-test, n= 12 per group.

## Discussion

The NR1^C2^ subunit was first identified almost 33 years ago^42^. By initially studying Magi-2 and using biochemical, electrophysiological, and behavioral approaches, we have identified the function of the NR1^C2^ subunit. It functions to confer susceptibility to ubiquitin ligases, answering a long-held question on the significance of this NR1 isoform^43^.

We found that Magi-2 deficiency was associated with a specific reduction in NR1^C2^ protein, whereas the other NMDAR subunits NR1^C2’^, NR2A, and NR2B were unchanged. We reasoned that ubiquitin ligase-mediated NR1^C2^ degradation was enhanced during Magi-2 deficiency based on our prior work with Magi-1 and Na_V_1.8 voltage-dependent sodium channels in dorsal root ganglion neurons ^14^. We reported that Magi-1 deficiency resulted in a loss of Na_V_1.8 protein and reduced inflammatory pain behavior ^14^ as Na_V_1.8 is susceptible to E3 ubiquitin ligase-mediated degradation^44^. Within the C-terminal of the NR1^C2^ subunit there is no Group I WW binding motif, but we identified a Group III WW binding domain. We subsequently demonstrated that NR1^C2^ is sensitive to Nedd4-1-dependent degradation. Therefore, NMDAR that contain NR1^C2^ are a labile pool of NMDAR with specific sensitivity to the Nedd4-1 UL. Indeed, it has been shown that in the basal state, NR1^C2^ abundance is, depending upon the brain region, anywhere from a fifth to a tenth of that of the alternatively spliced NR1^C2’^ ^45^. It is likely that the constant ubiquitin pressure maintains this ratio of NR1 C-terminal isoforms. During high activity states, this ratio may be altered affecting NMDAR trafficking and membrane expression. Several studies have shown NR1^C2^ to be dynamic, exhibiting altered levels of abundance following neuronal activity, but this was not shown for NR1^C2’38,46,47^.

Our study also reinforced the unique function of the Magi scaffold proteins: to protect proteins against UL-dependent degradation. PSD-95 remains the canonical post-synaptic density protein for excitatory synapses, principally involved in synaptic plasticity. For example, during nerve-injury induced-neuropathic pain, it was observed that thermal hyperalgesia, mechanical allodynia, and cold allodynia were all virtually absent in PSD-95 null mice. Although quite surprisingly, inflammatory pain behaviors, such as those produced by the CFA or formalin assay, were unaffected by PSD-95 deficiency^48^. Indeed, in Magi-2 deficient mice we observed no changes in AMPAR subunits or activity (Supplemental Fig. 5). These results would suggest that inflammatory pain signaling in the SCDH is dependent upon Magi-2 and NMDAR plasticity. Moreover, since Magi-2 is necessary for NR1^C2^ protein expression during UL degradative pressure, the Magi-2-NR1^C2^ interaction represents an attractive target for novel analgesics.

Our group has previously demonstrated the efficacy of peripherally administered lipidated peptides, designed to disrupt protein-protein interactions, in providing prolonged analgesia ^14,49^. The phosphorylation status of the mimetic peptides targeting the WW domain of Na_V_1.8 bidirectionally controlled pain behavior by modulating the degradation of these channels ^14^. Similarly, when we used the NR1^C2^ mimetic peptides to regulate pain behavior, the phosphorylation status of the threonine adjacent to the -LPR-motif bidirectionally modified CFA-induced pain behavior. This suggests that the amino acids flanking the WW binding domains are also relevant to a specific WW domain interaction. In unpublished data, we observed that the analgesic duration of an Na_V_1.8-WW targeted peptide lasts for 21-24 days (data not shown). This is consistent with a slow neuronal membrane phospholipid turnover ^50^ and lipidated peptide metabolism. Here we used a similar approach with lipidated NR1^C2^ derived peptides delivered by intrathecal injection and found that pain behavior can be bidirectionally regulated for many days after a single injection (Fig 5). The phosphorylated peptide provided robust analgesia with effects that are pain specific (Fig. 5), as contralateral measurements did not differ from baseline (Supplementary Fig. 13). We speculate that this phosphorylated peptide competes with NR1^C2^ containing NMDAR for Magi-2 binding exposing endogenous NR1^C2^ to ubiquitination and subsequent degradation. It remains to be seen whether native NMDAR containing the NR1^C2^ subunit are endogenously phosphorylated at T915. Surprisingly, a T915A mutated peptide was hyperalgesic for the duration of the CFA inflammatory pain assay. We speculate that the alanine-based peptide increases the stability of NR1^C2^ - containing NMDAR, protecting NR1^C2^ from ubiquitination and/or internalization during elevated neuronal activity. We did not observe an increase in NR1^C2^ levels in cultured hippocampal neurons treated with the alanine peptide. This is likely due to a lack of induced activity which may be necessary to drive the alanine peptide effect as was seen during pain. A basal level of ubiquitination pressure on NR1^C2^ at the membrane is likely present allowing phosphorylated peptide to decrease NR1^C2^ abundance in culture. We observed that hippocampal cultured neurons only express the Magi-2 beta isoform (Supplemental Figure 12). However, there may be a requirement for increased neuronal activity or increased signaling where the affinity of the alanine peptides for the WW-domains of Nedd4-1 is enhanced to protect NR1^C2^ from UL degradation in cultured neurons. It has been shown that Nedd4-1 activity is regulated by phosphorylation ^51^. It should be emphasized; the alanine peptide CFA pain behavioral data seems mainly driven by the effects in females. We have previously noted that CFA inflammatory pain in males and females differed in response to afferent activity manipulation; female inflammatory pain appears to have more of a central component ^52^. This data corroborates those findings and is consistent with the sex differences in pain observed in humans ^53,54^.

AMPAR lability in the context of synaptic plasticity is a well-documented phenomenon ^55^. The movement and turnover of AMPAR are orchestrated by NMDAR which are often thought of as being static during synaptic plasticity. In studies done in retinal ganglion cells, NMDAR, unlike AMPAR, were shown to reside outside the postsynaptic density^56^. An antibody that recognizes only the NR1^C2^ isoform was used in this study. Prior work on NMDAR surface dynamics at the neuronal plasma membrane have shown that NMDAR at the postsynaptic density exhibit confined diffusion with anchoring of receptors but high diffusion with low confinement in the extrasynaptic membrane^57^. These differing NMDAR pools, synaptic vs extrasynaptic, mobile vs immobile, might depend on whether NMDAR contain the NR1^C2’^ or NR1^C2^ subunit. As our data suggests, the C2 cassette appears to confer the ability of a subset of NMDAR to traffic in and out of the plasma membrane via Nedd4-1-dependent regulation. This NMDAR trafficking is highly relevant, because extrasynaptic NMDAR control synaptic currents and the intrinsic excitability of neurons^58^. Our study showed that this NMDAR trafficking can also be pharmacologically targeted opening novel therapeutic strategies to treat neurological disorders beyond pain, including neurogenerative diseases such as Alzheimer’s disease and psychiatric disorders such as schizophrenia.

## Acknowledgements

This work was supported by the National Institute of Health Grants NS113991 and NS128543. The authors would like to Thank Dr. Mark Baccei and Dr. Jie Li for their assistance with the spinal cord slice recording protocol. We would like to thank Dr. Wade Sigurdson of the University at Buffalo Confocal Microscopy Core Facility for his assistance with Confocal imaging of the spinal cord dorsal horn. We thank Dr. Elsa Daurignac for critical reading of this manuscript. Included illustrations were created using BioRender.com.

## Author Contributions

AB and GDS conceived the idea for the project and co-wrote the manuscript. GDS performed all electrophysiological, behavioral, and biochemical experiments, analyzed all the data, and generated the manuscript. MKM performed embryonic hippocampal dissections and neuronal cultures. AR conducted site-directed mutagenesis, and KH performed the baseline behavior measurements in Magi-2 floxed mice. All authors reviewed and provided critical feedback for the final manuscript.

## Competing Interests

AB is a co-founder of Channavix Therapeutics, LLC and Mimetic Medicines, INC. A provisional patent (serial number 63/598,137) has been filed on the use of lipidated peptidomimetics targeting NMDAR to treat pain. All other authors declare no competing interests.

## Materials and Methods

### Animals

All animal care, experiments, and procedures were conducted with approval from the University at Buffalo Institutional Animal Care and Use Committee (IACUC) in accordance with the guidelines set forth by the National Institutes of Health. Wild type C57Bl/6 mice were purchased from Envigo (Indianapolis, IN) and were used in Magi-2 viral knockdown and peptide experiments. Sperm from Magi-2 deficient mice were a kind gift from Dr. Katsuhiko Nishimori (Fukushima Medical University) and was sent to the Roswell Park Comprehensive Cancer Center Gene Targeting and Transgenic Shared Resource (Buffalo, NY) for *in vitro* fertilization of female WT C57Bl/6 mice. After breeding to obtain homozygous floxed mice, they were then backcrossed for several generations to obtain male and female mice heterozygotic for the floxed allele which were used for breeding in this study. Additionally, timed pregnant Sprague Dawley Rats used for embryonic hippocampal cultures were purchased from Envigo. Rats were singly caged on a 12:12 light dark schedule with food and water available *ad libitum*.

### Genotyping of Magi-2 fl/fl mice

A small segment of the tail was taken from pups at postnatal day 21 and stored at −20° C. Tails were then placed in a solution of 25mM Sodium Hydroxide/ 0.2mM EDTA ‘DNA extraction buffer’ and incubated at 98° C for 1hr. Following this, an equal volume of 40mM Tris-HCl (pH 5.5) was added to precipitate the gDNA, and the sample was placed on ice. Genomic DNA was PCR amplified using the following primers: forward 5’-AATAAAAATAGCTGTTTGAGGACAGGGAG-3’, reverse 1: 5’-GTCAAATAGAACCCACAGGGATGACAAAGA-3’, reverse 2: 5’-CATCGATTTTTTCCCAGCCATATGGAAGCT-3’ and resulting amplified DNA was visualized via gel electrophoresis (Fig S4b).

### Immunofluorescence

Wildtype C57Bl/6 mice were deeply anesthetized via intraperitoneal injection of 1mg/kg pentobarbital and then transcardially perfused with ice cold, filter-sterilized Phosphate Buffered Saline (PBS) followed by 4% Paraformaldehyde (PFA). Following fixation, lumbar spinal cord was removed and postfixed overnight in 4% PFA at 4° C and then allowed to equilibrate in sequential overnight incubations in 15% and then 30% sucrose at 4° C. Tissue was then embedded in Optimal Cutting Temperature (OCT) media, frozen, and stored at −80° C before slicing. 15-micron spinal cord slices were taken on a Leica CM1900 cryostat and attached to charged Superfrost microscope slides (Fisherbrand) and stored at −80° C. Prior to immunostaining, slides were thawed to room temp for 30 minutes. OCT media was then removed with a 5-minute wash in ice cold Tris-Buffered Saline (TBS) with 0.4% Triton-X-100. Antigen retrieval was then performed by incubating the slices in 0.3% w/v (pH 2.0 in ddH_2_O) Pepsin (Fischer, Nazareth, PA) at 37° C for 10 minutes. Slides were then washed 3X in 0.4% Triton-X-100 TBS for 5 minutes before blocking at room temperature for 2 hours in a solution of 10% Normal Goat Serum (Abcam), 3% BSA (Fisher) and 0.0.025% Triton-X-100 (Blocking Buffer). Following blocking, slides were quenched for 30 minutes with 10mM NH_4_Cl. Slides were next gently rocked in a humidity chamber overnight at 4° C with Blocking Buffer containing either Mouse anti-Magi-2 (1:250, Abnova), Guinea Pig anti-PKC*γ* (1:500, Nittobo Medical). Following this, slides were rinsed three times with 0.4% Triton-X-100 TBS and then incubated overnight with the secondary antibody. The secondary antibodies were diluted in blocking buffer as follows: Goat anti-mouse Alexa Fluor 488 (1:750, Abcam); Donkey anti-Guinea pig Alexa Fluor 594 (1:750, Abcam). Following the secondary antibody incubation, slides were rinsed three times for 5 minutes with 0.4% Triton-X-100 TBS and mounted using VectaShield® Hardset ™ Antifade Mounting Medium with DAPI (Vector Laboratories, California). Slides were then left alone to set for at least 48 hours at 4° C and imaged on a Leica DMi8 Confocal Microscope.

### Formalin Behavior

Male and Female WT, Heterozygous and ΔMagi-2 littermates were first acclimated to the testing room for a period of 30 minutes. Mice then received a unilateral intraplantar injection of 20*μ*L 5% Formalin diluted in PBS and were placed into a clear plexiglass recording chamber. Mice were video recorded using Active Webcam Software for 60 minutes following injection. After completion of all mice, videos were scored by blinded observers who counted the number of licks directed at- and lifts of- the injected hindpaw on separate views.

These behaviors were counted for 1-minute bins, every fifth minute for the total recording period of 60 minutes.

### Complete Freund’s Adjuvant Model of Inflammatory Pain

Baseline Thermal and Dynamic Weight Bearing Assessments were performed prior to all CFA injections. 20*μ*L of Imject® Complete Freund’s Adjuvant (Fisher) was administered via intraplantar injection into the right hindpaw of mice under isoflurane anesthesia. Mice were monitored following awakening from anesthesia and were left in their homecage until further behavioral testing was performed. We utilized the Imject® Complete Freund’s Adjuvant due to their storage in sealed glass ampules to ensure consistency of activity across experiments.

### Hargreaves Thermal Paw Withdrawal Latency

Mice were acclimated to the testing room first for a period of 15 minutes followed by placement in a clear plexiglass chamber on a raised platform for an additional 15-minute acclimation period. For testing of thermal paw withdrawal latency, an automatic Hargreaves Apparatus (Ugo Basil, Italy) was placed under the plantar surface and turned up to an intensity of 160mW/cm^2^ for a maximum of 15 seconds. Withdrawal latencies were calculated as the average of four applications to each hindpaw and each application was done with at least a five-minute interval.

### Dynamic Weight Bearing Assessment

For testing of weight distribution on the hind limbs, mice were first acclimated to the testing room for a period of 15 minutes and then placed within the Dynamic Weight bearing Apparatus (BIOSEB, France). The Dynamic Weight Bearing apparatus was placed behind a red plexiglass panel to ensure a uniform environment. Following a two-minute latency, mice were recorded for a period of 5 minutes during which time mice were able to freely roam the testing chamber which consisted of a platform made of small sensors that could measure the weight borne by each paw. For each acquisition, the BIOSEB DWB software matched each sensor activation to the paws and a blinded observer manually validated these paw assignments.

### Lumbar Spinal Cord Magi-2 Knockdown

Wildtype C57Bl/6 mice (Envigo) at 6 weeks of age were placed under isoflurane anesthesia and shaved over the lumbar section of the spinal cord. The spinal column was held in place with thumb and forefinger and the pelvic girdle was identified to locate the L3-L5 vertebrae. A small RN luer Hamilton syringe with 12.7mm, type-2 needle was then penetrated through the skin and placed between the vertebrae at an angle of ∼45° until resistance was felt. The angle of the syringe was then lowered to ∼30° and was then slid into the intrathecal space indicated by a tail flick response of the anesthetized mouse. 5 *μ*L of 2.0×10^13^ titer AAV2/9-GFP-U6-m-Magi-2-shRNA or AAV2/9-GFP-U6-m-scrmb-shRNA scrambled control (Custom, Vector BioSystems, Malvern, PA) was injected at a rate of about 0.1 *μ*L / sec and then held in place for an additional 15 seconds before removing the needle and placing the mouse back in its homecage for postsurgical observation. The sequence for the Magi-2 shRNA was based on prior findings ^34^. To confirm transduction of the lumbar dorsal horn, we performed immunohistochemical verification of GFP expression as described in ‘*Immunofluorescence’.* The antibodies used to probe for GFP was chicken anti-GFP (1:500, Abcam; ab13970) and Goat anti chicken Alexafluor 488 (1:750, Abcam; ab157169). The lumbar dorsal horn of a naïve, uninjeced mouse was used as a negative control. To verify knockdown of Magi-2, lumbar SCDH tissue was collected, and protein levels were determined as described in ‘*Ex-vivo* Western analysis’.

### *Ex-vivo* Western Analysis

Total protein was obtained from lumbar spinal cord dorsal horn tissue. Following dorsal laminectomy and excision of the spinal cord from the spinal column, the lumbar spinal cord was isolated and cut along the midline to segregate the right and left spinal cord. The hemisected spinal cord was then cut along the dorsal/ ventral axis to isolate and collect the dorsal horn. Tissue was homogenized in ice cold HEPES-buffered sucrose with protease inhibitors and centrifuged at 1000G for 15 minutes. The supernatant was then collected and re-centrifuged for 30 min at 10,000G to isolate the cytosolic (supernatant) and crude synaptic fraction (pellet). The pellet was resuspended in 1% SDS RIPA buffer and stored at −20° C. All samples were resolved on 4-20% Mini-PROTEAN TGX Precast Gels (BioRad) and transferred to 0.45um nitrocellulose membranes. The membranes were then blocked in 5% milk in Tris Buffered Saline with 1% Tween (TBS-T) overnight at 4° C before being probed overnight at 4° C with Rabbit anti Magi-2 (1:1000, Sigma M4221); Mouse anti-NR1 (1:1000, Abcam ab134308); Mouse anti-NR2A (1:1000, Abcam ab124913); Rabbit anti-NR2B (1:1000, Abcam ab254356); Rabbit anti-NR1-C2 (1:1000, PhosphoSolutions 1506-C2); Rabbit anti-NR1-C2’ (1:1000, PhosphoSolutions 1507-C2’); or Rabbit anti-actin (1:5000, Sigma a2066). Membranes were then washed 3x with TBS-T and incubated for 2 hours at room temperature with either anti-rabbit (1:5000, Promega W4011) or anti-mouse HRP (1:5000, PromegaW4021). Protein bands were visualized with Immobolin Forte Western HRP substrate (Sigma, WBLUF0100) and imaged with a ChemiDoc Imaging System (BioRad). Densitometric analysis was performed to quantify protein abundance relative to Actin using ImageJ software (NIH.gov). For all Western analysis experiments, each individual sample lysate was run in triplicate (three consecutive bands, technical replicates). The protein-to-actin ratio of the three bands was then averaged and represents a single sample contributing to the overall n-value for each experiment (biological replicates).

### Co-Immunoprecipitation

SCDH Lysates were collected as described for “*ex-vivo* Western Protein Analysis”. Following collection of the lysates and isolation of the crude synaptosomal fraction anti-Magi-2 antibody (Sigma) was added to half of the lysate and incubated at 4° C overnight with end over ending rocking. The next day, 50 *μ*L of Protein G Sepharose 4 Fast Flow affinity resin (Cytiva) was added to both lysates and incubated at 4° C for 3 hours.

Complexed protein was then isolated following centrifugation and rinsing followed by denaturation and resolution via SDS-PAGE. Immunoblotting for NR1 was performed as described for “*ex-vivo* Western Analysis”.

### Spinal Cord Slice Electrophysiology

Male and Female WT, Heterozygous, and ι1Magi-2 littermates aged 2-4 weeks were placed under deep anesthesia following an intraperitoneal injection of 1mg/kg pentobarbital. Mice were transcardially perfused with ice-cold cutting Ringer containing the following (in mM) 250 Sucrose, 2.5 KCl, 25 NaHCO_3_, 1 NaH_2_PO_4_, 6 MgCl_2_, 0.5 CaCl_2_, and 25 Glucose constantly bubbled with carbogen (95% O_2_, 5% CO_2_). Following laminectomy, the lumbar region of the spinal cord was isolated, and all dorsal and ventral roots were removed except for the right L3-L5 dorsal roots. Parasagittal slices (350um) with dorsal roots attached were taken using a LeicaVT1200S vibratome and transferred for 20 minutes into a recovery solution of the following contents (in mM) 92 N-methyl-D-glucamine, 2.5KCl, 1.2 NaH_2_PO_4_, 30 NaHCO_3_, 20 HEPES, 25 Glucose, 2 Sodium Ascorbate, 2 Thiourea, 3 Sodium Pyruvate, 10 MgSO_4_, and 0.5 CaCl_2_ bubbled constantly with carbogen (95% O_2_, 5% CO_2_). Slices were then allowed to recover for 1 hour in aCSF containing (in mM) 125 NaCl, 2.5 KCl, 25 NaHCO_3_, 1.0 NaH_2_PO_4_, 25 Glucose, 1 MgCl_2_, 2 CaCl_2_ bubbled constantly with carbogen (95% O_2_, 5% CO_2_). Slices were then transferred to a recording chamber (ALA Scientific) and visualized under a Zeiss Axioskop 2 FS Plus with a 40X water immersion lens equipped with DAGE MTI IR1000 differential contrast and interference system.

Patch electrodes were created from 4in thin wall borosilicate glass (TW150F-4, World Precision Instruments) and pulled with a Narishige Model PC-10 were filled with the following (in mM) 10.5 Cs-gluconate, 17.5 CsCl, 10 EGTA, 10 HEPES, 2MgATP, 0.5 Na_2_GTP, pH 7.25, 295 mOsm.

Whole-cell patch-clamp recordings were then taken from dorsal horn neurons that were located superficial to the translucent band delineating the substantia gelatinosa. Currents were amplified with a Multiclamp 700B amplifier and digitized with a Digidata 1440 Series. During recordings, slices were constantly bathed in room temperature CSF additionally containing 500nM Tetrodotoxin, 10 uM Cadmium, 10 uM Strychnine, and 10 uM Bicuculline to block voltage gated sodium channels, voltage gated calcium channels as well as glycine and GABA receptors, respectively. Neurons were held at −70mV to record spontaneous EPSCs for 1 min. After this, 10uM NBQX was added to the bath to block AMPA-mediated currents. Once spontaneous EPSCs were halted from bath application of NBQX, the neuron was slowly stepped up to +40mV. Membrane Current was recorded for one minute followed by bath application of 50uM NMDA for an additional 1 minute followed by a three-minute washout to ensure recovery. Neurons were then once again slowly stepped up to +60mV and recorded for one minute followed by a 1-minute bath application of 50uM NMDA and a three-minute washout. Only one neuron was recorded per slice to ensure there was no effect of bath applied NMDA prior to recording a neuron.

Currents were filtered at 5kHz and analyzed using pClamp10 software. Peak NMDA-induced currents recorded at +40mV and +60mV were taken and divided by the cell capacitance to determine total current density. Spontaneous EPCS current amplitudes and Frequencies were analyzed using Easy Electrophysiology Software (London).

### *In vitro* heterologous protein degradation assay and site-directed mutagenesis

Chinese Hamster Ovary cells cultured at 37° C with 5% CO_2_ in IMDM (Iscove’s Modified Dulbeccos Medium) containing 1% Hypoxanthine-Thymidine Supplement, 1% Penicillin-Streptomycin and 10% Fetal Bovine Serum. Cells were plated in six well plates and grown for 24 hours. CHO cells were then transiently transfected with 0.5 *μ*g of NR1^C2^, NR1^C2’^ or NR1^C2-P918A^ and 0.5 *μ*g Nedd4-1 or pcDNA using lipofectamine 2000. 48 hours later, cells were lysed in ice cold 1% RIPA Buffer containing protease Inhibitor cocktail. Samples were then frozen for western analysis of NR1 protein levels. Prior to western analysis, samples were mixed at a 1:1:1:1 ratio of sample: 8M Urea: 100uM DTT: 6X loading dye (0.06% w/v Tris-HCl, 0.06% w/v SDS, 0.05% v/v Glycerol, 0.1% v/v beta-mercaptoethanol, and bromoblue) in order to ensure proper denaturation and reduction of the NR1 subunit and ensure that single monomers were resolved via SDS-PAGE ^59^. NR1^C2^ and NR1^C2’^ plasmids were a gift from Dr. Gabriella Popescu, SUNY-Buffalo. The NR1^C2-P918A^ was created via site-directed mutagenesis using the QuickChange Site-directed Mutagenesis Kit (Agilent Technologies). NR1^C2^ plasmid and mutation primers were reacted with PfU turbo and then digested with Dpn1. The primer sequences for the mutation primers are F: 3’-caaaaagacacagtgctgctggcgcgacgcgctattgag-5 and R: 3’-cacaatagcgcgtcgcgccagcagcactgtgtctttttg-5’. Resulting PCR products were then transformed into One-Shot Top Ten Competent cells (ThermoFischer) and cultured overnight. Plasmid DNA was extracted using the Qiagen Maxi-prep kit and frozen in aliquots for later use in transient transfection experiments. All plasmids used underwent full Plasmid sequencing through Plasmidsaurus to confirm identity (Eugene OR). Sequences available upon request.

### Cultured Hippocampal Neurons

Timed-pregnant Sprague Dawley Rats (Envigo; Indianapolis, IN) were used for embryonic hippocampal cultures. The timed pregnant rat was sacrificed on day E18 by CO_2_ asphyxiation and the embryos were extracted. After identification and extraction of the hippocampi in ice-cold dissection solution consisting of 7.5mM NaHCO_3_, 10mM HEPES, 100 U/mL Penicillin/Streptomycin (Gibco) in Hanks Balanced Salt Solution (Corning) adjusted to 310mOsm with sucrose, neurons were digested in 0.5% trypsin for 20 minutes at 37°C with constant agitation. Next, the neurons were plated on laminin-coated plates with Plating Media consisting of 1mM Sodium Pyruvate, 0.6% Glucose, 10% Horse Serum (Gibco), 2mM Glutamax (Gibco), and 100 U/mL Penicillin/Streptomycin (Gibco) in Minimum Essential Medium (MEM) (Corning). The cultures were maintained at 37°C in a humidified incubator with 7% CO_2_ overnight before replacing the plating media with growing media the day after the dissection. Growth Media consisted of 1% B27 (Gibco), 2mM Glutamax (Gibco), 100 U/mL Penicillin/Streptomycin (Gibco) in Neurobasal A (Gibco). Subsequently, half of the media was replaced for the first 5 days after dissection, with the addition of 3μM cytosine-Δ-D-arabinofuranoside hydrochloride (Sigma) on days 3 and 4. After day 5 half of the media was replaced every other day. Neurons were treated with 10μM peptides added to the growth media DIV6 and incubated overnight. On DIV7, neurons were washed twice with ice-cold PBS before lysis in ice-cold 1% SDS RIPA buffer containing protease inhibitor cocktail (Sigma). Lysates were centrifuged at 12,000G for one hour and the supernatant was collected and frozen at −20° C until further use. Lysates were then processed as described in ‘*Ex-vivo* Western Analysis’.

### *in-vivo* Intrathecal Peptide Injections

Male and Female WT C57Bl/6 mice were first tested for baseline thermal withdrawal latency and dynamic weight bearing hind limb distribution for two days. Immediately after behavioral testing on the second day, mice were injected with 20 *μ*L of CFA into the right hindpaw under isoflurane anesthesia. Twenty-four hours later, thermal withdrawal latency and dynamic weight bearing was assessed. After behavioral testing mice were placed under isoflurane anesthesia and shaved over the lumbar section of their spinal cord. As with the intrathecal virus injections, the spinal column was held in place with thumb and forefinger and the pelvic girdle was identified to locate the L3-L5 vertebrae a small RN luer Hamilton syringe with 12.7mm, type-2 needle was then penetrated through the skin and placed between the vertebrae at an angle of ∼45° until resistance was felt. The angle of the syringe was then lowered to ∼30° and was then slid into the intrathecal space. N-terminal-myristoylated peptides based were on based on the C-terminal sequence of the mouse NR1^C2^ and custom ordered from GenScript. The wildtype sequence was NQKDTVLPRRAIERE. The phosphorylated peptide was NQKDT^P^ VLPRRAIERE and the alanine replacement peptide was NQKDAVLPRRAIERE. The scrambled peptide sequence was NARLIRDEQKTVRPE. All peptides were made up as previously described ^52^. 5 *μ*L of 100 *μ*M peptide solution dissolved in 0.2% DMSO in PBS was then injected slowly at a rate of about 0.1 *μ*L per second. Once the total volume was injected, the needle was held in place for an additional 15 seconds and removed. The mice were then observed post-operatively and tested behaviorally again at 48 and 72 hours relative to the CFA injection and again on days 5,7, 10, and 14. A total of 12 mice (6 male and 6 female) were tested per peptide.

### Statistics

All statistical analyses were performed using GraphPad Prism Version 9. All data is represented as mean +/- SEM. Statistical significance was determined by setting a p-value threshold of p<0.05 for all experiments. Simple between-group comparisons were done using unpaired students t-test where appropriate. All multiple comparisons were analyzed via one- or two-way ANOVA with Tukey post hoc test or Bonferroni correction, respectively for comparison between groups. pClamp10 software, Easy Electrophysiology and Origin 8 were used to analyze all electrophysiology data. Densitometric analysis was performed using ImageJ software.

---

**Supplemental Figure 1: Magi-2 isoform ratio in spinal cord dorsal horn lysates of WT C57Bl/6j mice.** Spinal cord dorsal horn tissue lysates were harvested from 8-week-old WT mice and processed through differential centrifugation (see materials and methods) to acquire the crude synaptosomal and cytosolic fractions. **A & B**) As previously reported, ^9^ only the Alpha and Beta isoforms were present in the cytosolic fraction, while all three isoforms, including the gamma isoform was present in the synaptic fraction. This suggest that the gamma isoform is locally translated. **C & D**) Parts of whole representation of the isoform distribution in each fraction. n=6, 3 males and 3 females.

**Supplemental Figure 2: Immunocytochemical verification of Mouse monoclonal antibody raised against Magi-2** Chinese Hamster Ovary Cells were cultured and transiently transfected using Lipofectamine Reagent with plasmid DNA carrying Magi-2. Lipofectamine transfection gives ∼20% transfection efficiency. 48 hours after transfection, cells were fixed and stained with mouse monoclonal Magi-2 antibody (Abnova) 1:250 (left) followed by goat ant-mouse Alexafluor 488 (1:750) (left and right) and mounted with Vectashield Hardset mounting media with DAPI. Images were acquired using a Leica DMi8 equipped with LasX acquisition software. Scale Bar is 66.5um.

**Supplemental Figure 3: Immunofluorescent staining using a polyclonal Magi-2 antibody**. Immunolabeling using this antibody similarly shows labeling in the superficial dorsal horn, recapitulating staining seen with monoclonal mouse Magi-2 antibody (Fig1b). Spinal cord sections were obtained as described in material and methods. Co-staining was performed to determine colocalization relative to PKC*γ* and TACR1

**Supplemental Figure 4: Further Characterization of the floxed ΔMagi-2 mouse line** (**A**) The genetic loxP insertion strategy to generate Magi-2 floxed mouse (adapted from Shirata et al ^13^). The endogenous 3’ UTR is altered in this genetic strategy perhaps explaining the Magi-2 deficiency **B**) Image of agarose gel showing PCR product for genotyping of ΔMagi-2 mice. Primer sequences available in Materials and Methods. **C**) Western analysis showing the cytosolic fraction of lysates described in Fig 1C. **D**) Quantification by densitometric analysis of Magi-2 Alpha and Beta levels in cytosolic fraction demonstrated in C. **E**) Baseline thermal withdrawal latencies showed no differences across genotypes using Hargreaves apparatus. **F**) Baseline mechanical withdrawal thresholds measured via ascending von Frey Fibers showed no differences across genotypes

**Supplemental Figure 5: No differences in spontaneous EPSC (sEPSC) properties or synaptic AMPA receptor protein levels across genotypes of Magi-2 deficient mice A**). Whole-cell patch-clamp recordings were obtained from lamina 1 neurons as described in Materials and Methods. Prior to bath application of NMDA, neurons were held at −70mV for one minute to obtain baseline current recordings. This timeframe was then analyzed using Easy Electrophysiology software to obtain the average amplitude, frequency and decay fit tau values for sEPSC. **B-D**) Events were detected with a threshold of -5pA. **E**) Western analysis of AMPA receptor subunits GluA1 and GluA2 from the synaptic fraction of spinal cord dorsal horn lysates. These were the same lysates used to determine Magi-2 deficiency in Fig 1 and Sup Fig 3. **F**) Densitometric analysis revealed no differences in the levels of GluA1 or GluA2 across genotypes. One-way ANOVA with Tukey Post hoc test for all comparisons across genotypes

**Supplemental Figure 6: Confirmation of SCDH viral transduction and densitometric analysis of the western analysis from** Fig 3b **separated by isoform**. (Top) Immunohistochemical verification of GFP expression in the lumbar dorsal horn two weeks after intrathecal injection of AAV2/9-GFP-U6-m-Magi-2-shRNA using an antibody against GFP (right) versus a naïve mouse stained and imaged under the same conditions (left); scale bar is 100um. (Bottom) Alpha: shRNA VS. Scram p=0.0115*; Beta: shRNA VS. Scram p=0.0059**; Gamma: shRNA VS. Scram p=0.0033**. Unpaired students t-test: n=8 scram, n=6 shRNA

**Supplemental Figure 7: Baseline thermal paw withdrawal latencies and segregation by sex during Magi-2 knockdown**. **A**) Baseline thermal paw withdrawal latencies show no differences across groups prior to either intrathecal administration of AAV2/9-GFP-U6-m-Magi-2-shRNA or AAV2/9-GFP-U6-m-Magi-2-scram control. **B**) Thermal Paw Withdrawal Latencies for only the male cohort: shRNA vs. Scram; 24 hours post-CFA: p=0.0028**; Day 3: p=0.0121*; Day 5 p=0.0123* **C**) Thermal Paw withdrawal latencies for only the female cohort: shRNA vs Scram: 3 days post-CFA: p=0.0016**; Day 5: p=0.0293*, Day 10: p=0.0297*; Day 14: p=0.0066**.**D**) direct comparison of males vs. females revealed no sex differences. All data represented as mean PWL +/- SEM, Two-way ANOVA with Bonferroni Correction n=11 scram (5 males, 6 females) n=12 shRNA (6 males, 6 females).

**Supplemental Figure 8: NR1^C^**^2^ **is selectively lost in ΔMagi-2 mice. A**) Western analysis of NR1^C2^ in WT, Heterozygous and ΔMagi-2 mice. **B**) Densitometric Analysis of A: WT vs. Het p=0.0201*. **C**) Western analysis of NR1^C2’^ in WT, Heterozygous and ΔMagi-2 mice. **D**) Densitometric Analysis of C. No differences were observed. Data represented as mean protein/ actin. One-Way ANOVA with Tukey post hoc test. Each set of three bands is from a single lysate, the density of these three bands relative to the actin for each lane from each lysate are then averaged to create an n of 1 from a single animal. n=5WT, n=5Het, n=5ΔMagi-2

**Supplemental Figure 9: Sequence alignment of NR1 C2 and C2’ C-termini from various species.** Red denotes PDZ binding domain. Blue denotes Group III WW binding domain. NR1^C2^ is only found in mammals.

**Supplemental Figure 10: Proteasome inhibitor MG-132 blocks the ability of Nedd4-1 to promote the degradation of NR1^C^**^2^**. A**) Western analysis showing the effect of MG-132 (50uM) treatment on NR1^C2^ levels. **B**) Western analysis showing the effect of MG-132 (50uM) treatment on NR1^C2^ levels co-transfected with Nedd4-1. **C**) Densitometric analysis of A reveals no difference in NR1^C2^ levels upon treatment with MG-132. **D**) Densitometric Analysis of B shows that MG-132 blocks the Nedd4-1 dependent loss of NR1^C2^ levels. Vehicle (0.1% DMSO in PBS) vs. MG-132 p= 0.0498*, paired students t-test n=3. Data represented as mean protein level relative to actin +/- SEM

**Supplemental Figure 11: NR2B is selectively susceptible to Nedd4-2 but not Nedd4-1 mediated degradation while NR2A unaffected by either**. **A**) Representative western analysis for NR2A following co-transfection with Nedd4-1. **B**) Representative western analysis for NR2A following co-transfection with Nedd4-2. **C**) Densitometric analysis for A showing no difference in NR2A protein levels upon co-transfection with Nedd4-1. **D**) Densitometric analysis for A showing no difference in NR2A protein levels upon co-transfection with Nedd4-2. **E**) Representative western analysis for NR2B following co-transfection with Nedd4-1. **F**) Representative western analysis for NR2B following co-transfection with Nedd4-2. **G**) Densitometric analysis for E showing no difference in NR2B protein levels upon co-transfection with Nedd4-1. **H**) Densitometric analysis for F: pcDNA vs. Nedd4-2 p=0.0467*. paired students t-test n=3/group

**Supplemental Figure 12: Western analysis to probe for presence of Magi-2 in cultured hippocampal neurons**, a A rabbit polyclonal antibody was used that recognizes all three isoforms of Magi-2. Only the beta form was detected in hippocampal cultures

**Supplemental Figure 13: Contralateral paw withdrawal latencies of peptide and CFA treated mice** from Main Text Figure 5B.

